# RAB23 Modulates Signaling Protein Ciliary Homeostasis through Promoting RAB18-mediated Inward BBSome Transition Zone Passage

**DOI:** 10.1101/2025.02.20.639385

**Authors:** Sheng-Nan Sun, Wen-Jian Sun, Yan Wei, Zhen-Chuan Fan

## Abstract

As an intraflagellar transport (IFT) cargo adaptor, the BBSome targets the basal body region located just below the transition zone (TZ) before proceeding to cross the TZ for ciliary entry. This process of TZ crossing crucially governs BBSome presence and quantity available for signaling protein coupling within cilia, influencing the dynamic behavior of signaling proteins in cilia by controlling their ciliary removal. Our previous study revealed that the GTPase RAB18 binds to its BBSome effector in the basal bodies right below the TZ and recruits the BBSome to cross the TZ for ciliary entry by lateral transport between the plasma and ciliary membranes. However, the mechanism by which RAB18 is activated to accomplish this function has remained elusive. This study uncovers that, under physiological conditions, *Chlamydomonas* RAB23 in its GDP-bound state (RAB23-GDP) predominantly localizes to the same basal body region as RAB18-GDP and activates RAB18-GDP as a RAB18-specific guanine nucleotide exchange factor. RAB18-GDP attaches to the membrane and binds to the BBSome. This interaction occurs independently of anterograde IFT association, allowing the BBSome to diffuse through the TZ and enter cilia. Subsequently, RAB23-GDP facilitates its own ciliary entry by binding to the RAB18-GTP/BBSome entity as a BBSome cargo. Our findings establish a model where the BBSome enters cilia through the RAB23-RAB18 module-mediated inward BBSome TZ diffusion pathway. According to this model, hedgehog signaling defects caused by *rab23* mutations could induce Carpenter syndrome in humans, providing a mechanistic understanding behind the inward BBSome TZ passage required for proper ciliary signaling.

**Significance statement:** Certain ciliary signaling proteins are removed from cilia through the outward BBSome transition zone (TZ) diffusion pathway. The inward BBSome TZ crossing is orchestrated by RAB18. The current study unveils that RAB23 in its GDP-bound state (RAB23-GDP) and RAB18-GDP predominantly localize to the same basal body region located just below the TZ. RAB23-GDP activates RAB18-GDP as a RAB18-specific guanine nucleotide exchange factor. RAB18-GTP subsequently recruits the BBSome as its effector to cross the TZ for ciliary entry. Following this, RAB23-GDP enters cilia as a BBSome cargo. The regulatory interplay between RAB23 and RAB18, with RAB23 promoting inward BBSome TZ crossing via RAB18, establishes a molecular mechanism for the BBSome-dependent retrieval of signaling proteins from cilia.

## Introduction

Cilia project from the surface of eukaryotic cells, functioning as sensory antennae that detect and transduce extracellular stimuli into intracellular responses (1, 2). To ensure proper functionality, membrane-anchored sensory proteins, including G protein-coupled receptors and ion channels, along with their downstream signaling factors, must maintain their dynamics in cilia by cycling between cytoplasma and cilia (3). Within cilia, these proteins act as cargoes for intraflagellar transport (IFT) trains composed of repeated complexes IFT-A and -B (subdivided into IFT-B1 and -B2) (4–6). The BBSome, consisting of multiple Bardet-Biedl syndrome (BBS) proteins, serves as an IFT cargo adaptor, connecting sensory proteins to IFT (7–10). Across species, the presence and quantity of the BBSome in cilia are tightly regulated during various BBSome ciliary cycling steps, including targeting to the basal bodies (9, 11), diffusing through the ciliary diffusion barrier, the transition zone (TZ), for ciliary entry (12), ciliary tip remodeling and turnaround (13, 14), and diffusing through the TZ for ciliary retrieval (15, 16). Disruptions in these BBSome ciliary cycling steps lead to the loss/retention of sensory proteins in cilia (7, 8, 10, 17–19). This impairs proper ciliary signaling and resulting in the ciliopathic disorder BBS (20, 21).

During BBSome ciliary cycling, the Rab-like 5 (RABL5) GTPase IFT22 binds the Arf-like 6 (ARL6) GTPase BBS3 to form a heterodimer (BBS3/IFT22) in the cell body (11). Only in their GTP-bound states does BBS3/IFT22 associate with the BBSome and recruit it to the basal bodies, a critical step in controlling BBSome availability for ciliary entry (11, 22). To prevent competition with BBS3/IFT22 for BBSome binding in the cell body, the Arf-like (ARL) GTPases ARL13 and ARL3 target the basal bodies in their GDP-bound states (ARL13-GTP and ARL3-GTP) before entering cilia (16, 23). Upon ciliary entry, nucleotide exchange occurs, facilitated by ARL6/BBS3 and RABL2 acting as guanine nucleotide exchange factors (GEFs) for ARL13 and ARL3, respectively (15, 16, 23). ARL13-GTP recruits its BBSome effector to anchor to the ciliary tip membrane, allowing coupling of the BBSome with signaling proteins (23, 24). ARL3-GTP, by contrast, binds its signaling protein-laden BBSome effector in the proximal ciliary base localized right above the TZ (16). This interaction facilitates the cargo-laden BBSome to diffuse through the TZ for ciliary retrieval via lateral transport between the ciliary and plasma membranes (16, 25). Similarly, the Ras-associated binding (RAB) GTPase RAB18, in its GDP-bound state, targets the basal bodies, where it undergoes activation to RAB18-GTP (12). Once activated, RAB18-GTP binds its BBSome effector, promoting its diffusion through the TZ for ciliary entry via RAB18-mediated lateral transport between the plasma and ciliary membranes (12). Despite these insights, the molecular mechanism underlying RAB18-GDP activation remains to be elucidated.

Rab23 has been shown to regulate the ciliary levels and transport of Smoothened, which correlates with Rab23’s role as a sonic hedgehog (Shh) signaling inhibitor (26–30). This study identifies RAB23-GDP is enriched at the basal bodies. In this compartment, RAB23-GDP functions as a RAB18-specific GEF, catalyzing the activation of RAB18-GDP to RAB18-GTP. This activation facilitates the BBSome to diffuse through the TZ for ciliary entry via the RAB23-RAB18 module (12). Following RAB18 activation, RAB23-GDP dissociates from RAB18-GTP and subsequently associates with the RAB18-GTP/BBSome complex as a BBSome cargo, enabling its own diffusion through the TZ for ciliary entry. This study addresses a significant gap in understanding how the RAB23-RAB18 module promotes inward BBSome TZ crossing for ciliary entry. It suggests that deficiencies in RAB23 could disrupt this process, potentially leading to BBS-like phenotypes in humans.

## Results

### RAB23 diffuses into cilia without affecting IFT

*Chlamydomonas* RAB23 shares approximately 50% sequence homology with its orthologues in other ciliated species and is phylogenetically more distantly related to mammals and humans (*SI Appendix,* Fig. S1*A*-*C*). Using our newly developed RAB23 antibody, RAB23 was confirmed to localize to cilia, consistent with previous ciliary proteomics analysis of *C. reinhardtii* (Fig. 1*A*, *SI Appendix,* Fig. S2*A*) (31). Attempts to use the CLiP strain (LMJ.RY0402.185544, referred to as *rab23-544*), which contains an insertion at the 5’-proximal region of the first exon of the *rab23* gene was unsuccessful, as this strain was not an authentic RAB23-null mutant (*SI Appendix,* Fig. S2*B*-*D*). Efforts to generate a RAB23-null mutant via the CRISPR/Cas9 system also failed (32, 33). We instead utilize a highly efficient artificial miRNA system to silence RAB23 expression (34). A knockdown (KD) strain we generated, named *RAB23^miR^* was derived from the CC-5325 genetic background and showed a 91% reduction in RAB23 protein levels compared to parental CC-5325 cells, as determined by immunoblotting of whole cell samples (Fig. 1*A*). *RAB23^miR^* cells retained normal levels of IFT-A (IFT43), IFT-B1 (IFT22), and IFT-B2 (IFT38) subunits in both whole cell samples and cilia, excluding a role for RAB23 in a steady-state IFT (Fig. 1*A*). Furthermore, *RAB23^miR^* cells exhibited normal cilium growth, suggesting that RAB23 is not essential for ciliation in *C. reinhardtii* (Fig. 1*B*, *SI Appendix,* Fig. S3*A* and *B*).

**Figure 1.**
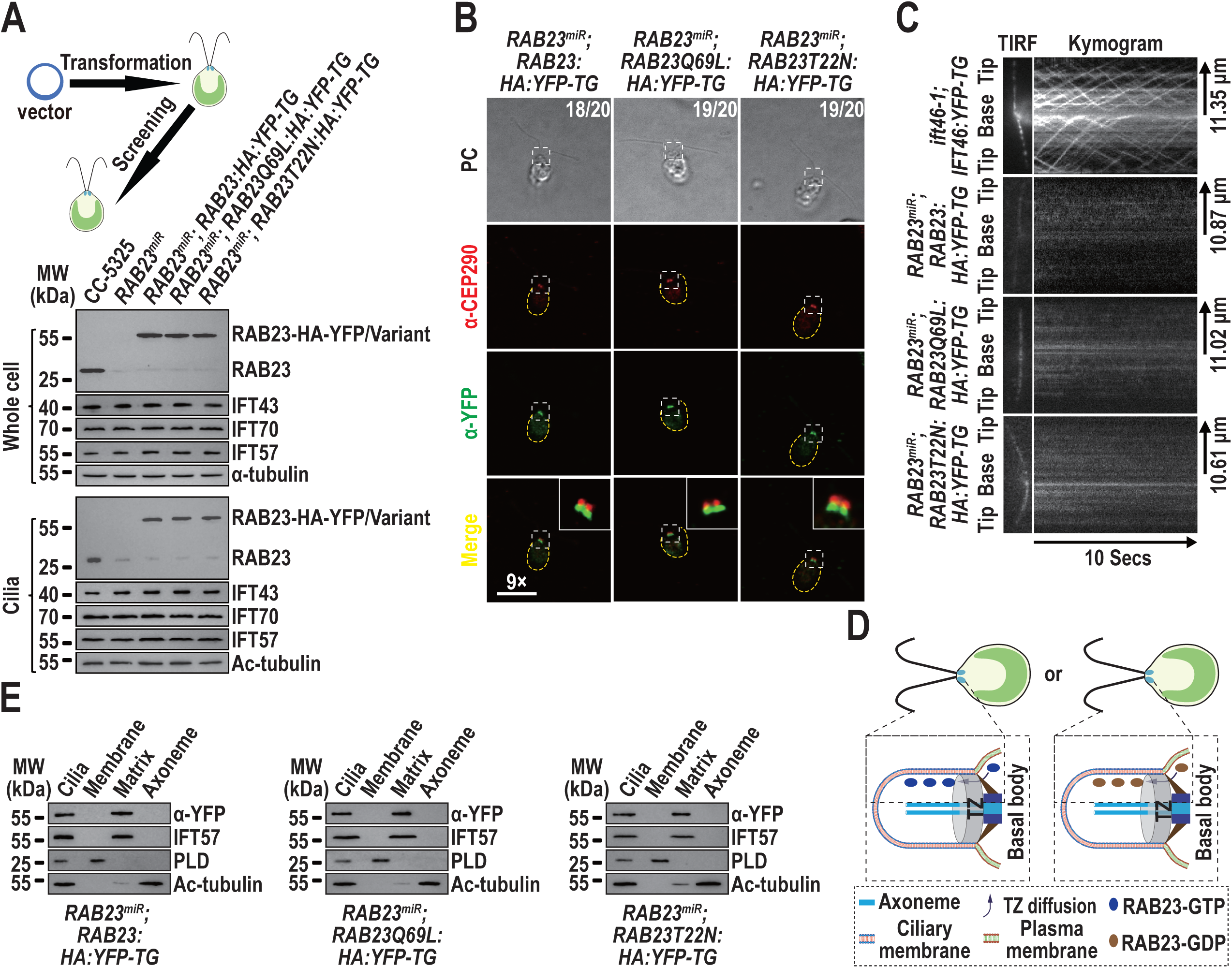
RAB23 diffuses into cilia without affecting IFT. (*A*). Immunoblots of whole cell samples and cilia from the indicated strains (top) probed with α-RAB23, α-IFT43 (IFT-A), α-IFT57 (IFT-B1), and α-IFT70 (IFT-B2). Alpha-tubulin and acetylated (Ac)-tubulin served as loading controls for whole cell samples and cilia, respectively. MW indicates molecular weight. (*B*) Immunofluorescence staining of the indicated cells (top) with α-CEP290 (red) and α-YFP (green). Representative phase contrast (PC) images of cells are shown. Insets displays 9 times magnification of the basal body and the TZ, marked by the gray box. Scale bar: 10 µm. Cell counts (out of 20) are listed for HA-YFP-tagged RAB23, RAB23Q69L, or RAB23T22N observed in the basal body region below the CEP290-labeled TZ. (*C*). Representative TIRF images and corresponding kymograms of the indicated cells (left) (Videos S1-S4, 15 fps). The time (bottom) and transport lengths (right) are indicated. The ciliary base (base) and tip (tip) are labeled. (*D*). Schematic representation of the diffusion of RAB23 in different nucleotide state into cilia via TZ passage. (*E*). Immunoblots of ciliary fractions from the indicated strains (bottom) probed with α-YFP, α-IFT57 (ciliary matrix marker), α-PLD (ciliary membrane marker), and α-Ac-tubulin (axoneme marker). MW indicates molecular weight.

To investigate the nucleotide state in which RAB23 enters cilia, we restored RAB23 expression in *RAB23^miR^* cells by expressing RAB23 fused at its C-terminus to hemagglutinin (HA) and yellow fluorescent protein (YFP) (RAB23-HA-YFP), as well as two mutant forms, RAB23Q69L-HA-YFP, and RAB23T22N-HA-YFP (*SI Appendix,* Fig. S1*A*). These resulted in the strains *RAB23^miR^; RAB23:HA:YFP-TG, RAAB23^miR^; RAB23Q69L:HA:YFP-TG*, and *RAB23^miR^; RAB23T22N:HA:YFP-TG*). These strains expressed RAB23-HA-YFP in a wild-type, active (Q69L, GTP-bound state), and inactive (T22N, GDP-bound state), respectively, together with the trace amount of RAB23, at the endogenous RAB23 level of the parental CC-5325 cells (Fig. 1*A*). Neither mutation impaired the ability of RAB23-HA-YFP to enter cilia and both localized at levels predominantly exceeding those of the endogenous protein within cilia in *RAB23^miR^* cells (Fig. 1*A*). Both mutant proteins resembled RAB23-HA-YFP in localizing to the basal bodies located right below the CEP290-labeled TZ prior to ciliary entry (Fig. 1*B*). As the endogenous RAB23 exhibited the same basal body localization pattern as the tagged proteins, the tags at the C-termini unlikely interfere with RAB23 intracellular trafficking and function (*SI Appendix,* Fig. S4). Using total internal reflection fluorescence (TIRF) microscopy, we observed that neither RAB23Q69L-HA-YFP nor RAB23T22N-HA-YFP underwent IFT as shown by their lack of movement compared to IFT46-YFP in the *ift46-1; IFT46:YFP-TG* strain (Fig. 1*C*, Videos S1-S4) (35). Instead, both entered cilia via diffusion (Fig. 1*C*, Videos S1-S4). Within cilia, both mutants, like RAB23-HA-YFP, were distributed along the entire ciliary length (Fig. 1*C*). These results suggest that RAB23, regardless of its nucleotide state, can enter cilia via diffusion through the TZ (Fig. 1*D*). Once inside cilia, it resided in the ciliary matrix, as both RAB23Q69L-HA-YFP and RAB23T22N-HA-YFP were detected exclusively in the ciliary matrix fraction (Fig. 1*E*).

### RAB23-GDP and the BBSome exhibit interdependence during inward TZ passage

Given that RAB23 enters cilia via diffusion without disrupting IFT, we ought to understand how RAB23 might regulate BBSome homeostasis within cilia. To explore this, we examined the RAB23 KD strain *RAB23^miR^* and the corresponding rescue strains *RAB23^miR^; RAB23:HA:YFP-TG*, *RAB18^miR^; RAB23Q69L:HA:YFP-TG* and *RAB23^miR^; RAB23T22N:HA:YFP-TG*. Notably, even in the complete absence of RAB23, the BBSome maintained normal cellular levels (Fig. 2*A*) and effectively targeted the basal bodies (Fig. 2*B*). However, BBSome entry into cilia relied on the presence of RAB23 in its GDP-bound state (Fig. 2*A*). To investigate the BBSome’s localization relative to the TZ before ciliary entry, we significantly reduced RAB23 expression (by about 95%) in the *bbs8; BBS8:YFP-TG* strain, resulting in the strain *RAB23^miR^; bbs8; BBS8:YFP-TG*. This strain expresses BBS8-YFP at the endogenous BBS8 level on a BBS8-null mutant *bbs8* background (Fig. 2*C*) (16). As expected, the absence of RAB23 impeded the BBSome (BBS8-YFP) entry into cilia (Fig. 2*C*), although it (represented by BBS8-YFP) successfully targeted the basal bodies right below the CEP290-labeled TZ (Fig. 2*D*). These findings underscore the dependence of the BBSome on RAB23 for its passage across the TZ and entry into cilia. To dissect the reciprocal interaction between RAB23 and the BBSome during inward TZ passage into cilia, we generated the strains *RAB23^miR^; bbs8; RAB23:HA:YFP-TG*, *RAB23^miR^; bbs8; RAB23Q69L:HA:YFP-TG*, and *RAB23^miR^; bbs8; RAB23T22N:HA:YFP-TG*. These strains expressed RAB23-HA-YFP or its variants at levels comparable to endogenous RAB23 in a RAB23 KD and BBS8-null mutant background (Fig. 2*E*) (16). The absence of BBS8 disrupts BBSome assembly in the cytoplasm, devoid of an intact BBSome available for targeting the basal bodies (IFT81-labeled) for ciliary entry (Fig. 2*E* and *SI Appendix,* Fig. S5*A-B*) (36). Strikingly, the absence of the BBSome completely prevented both RAB23-HA-YFP and its T22N mutant from entering cilia (Fig. 2*E*), although they correctly localized to the basal bodies localized right below the CEP290-labeled TZ (Fig. 2*F*). Conversely, the Q69L mutant targeted the basal bodies right below the CEP290-labeled TZ (Fig. 2*F*) and entered cilia in the absence of the BBSome (Fig. 2*E*). Since RAB23-HA-YFP resembles its T22N mutant version in targeting the basal bodies and failing to enter cilia, we conclude that RAB23 must exist in a GDP-bound state at the basal bodies under physiological conditions. RAB23-GDP and the BBSome rely on each other for inward TZ passage and ciliary entry.

**Figure 2.**
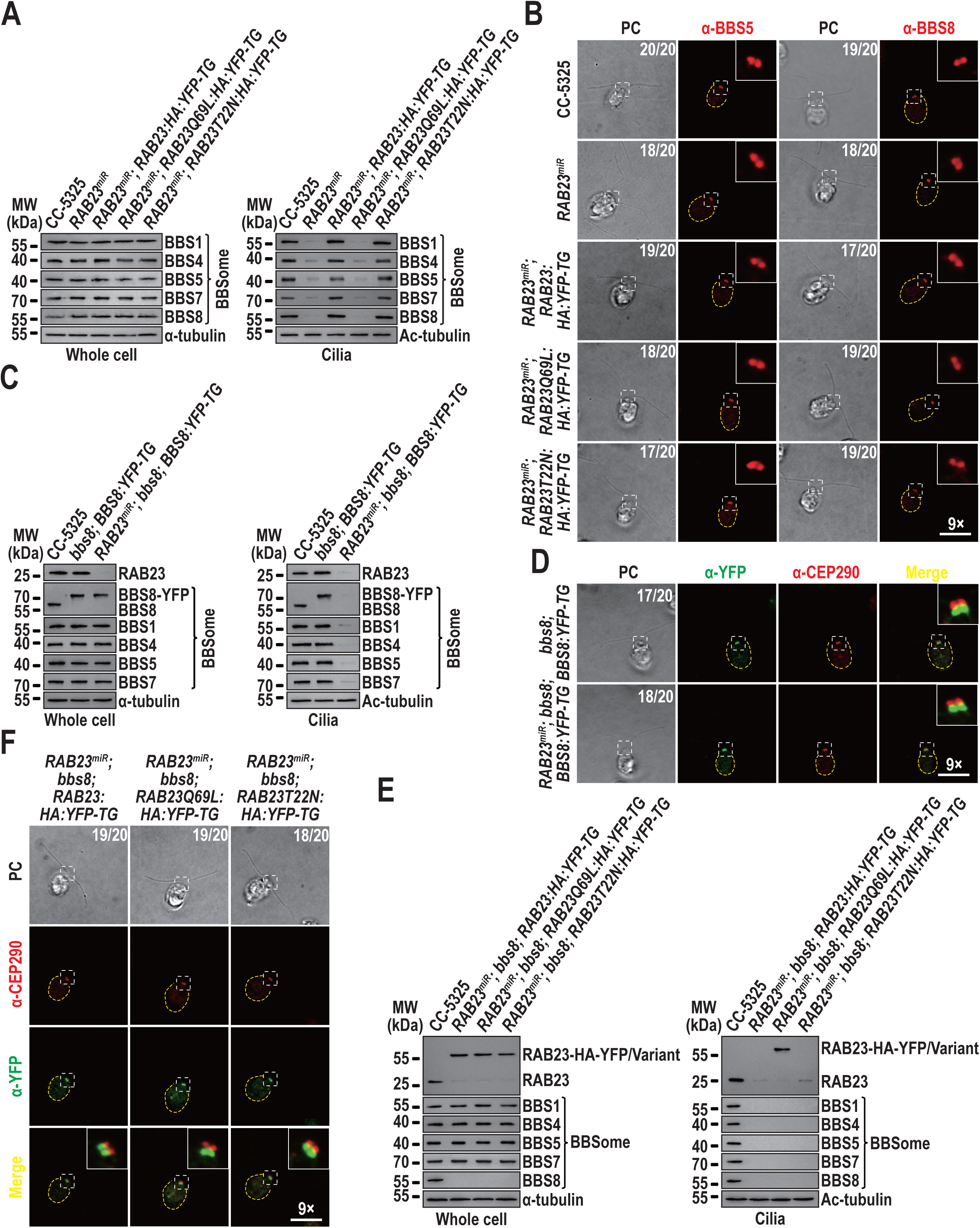
RAB23-GDP and the BBSome rely on one another for inward TZ passage. (*A*). Immunoblots of whole cell samples and cilia from the indicated cells (top) probed with α-BBS1, α-BBS4, α-BBS5, α-BBS7, and α-BBS8. (*B*). Immunofluorescence staining of the indicated cells (left) with α-BBS5 and α-BBS8. Cell counts (out of 20) represent cells positive for BBS5 or BBS8 at the basal bodes. (*C*). Immunoblots of whole cell samples and cilia of the indicated cells (top) probed with α-RAB23, α-BBS8, α-BBS1, α-BBS4, α-BBS5, and α-BBS7. (*D*). Immunofluorescence staining of the indicated cells (left) with α-CEP290 (red) and α-YFP (green). Cell counts (out of 20) represent cells positive for BBS8-YFP and CEP290 at the basal bodes and the TZ. (*E*). Immunoblots of whole cell samples and cilia from the indicated cells (top) probed with α-RAB23, α-BBS1, α-BBS4, α-BBS5, α-BBS7, and α-BBS8. (*F*). Immunofluorescence staining of the indicated cells (top) with α-CEP290 (red) and α-YFP (green). Cell counts (out of 20) represent cells positive for BBS-YFP and CEP290 at the basal bodes and the TZ. For panels *A*, *C*, and *E*, α-tubulin and Ac-tubulin serve as loading controls for whole cell samples and cilia, respectively. MW indicates molecular weight. For panels *B*, *D*, and *F*, representative PC images are displayed. Insets displays 9 times magnification of the basal body and the TZ, marked by the gray box. Scale bar: 10 µm.

### RAB23-GDP facilitates inward BBSome TZ passage by activating RAB18

Our previous study identified RAB18 in a GTP-bound state binds its BBSome effector, independent of association with anterograde IFT trains, at the basal bodies (12). By anchoring to the plasma membrane, RAB18-GTP enables BBSome diffusion across the TZ for ciliary entry via lateral transport between the plasma and ciliary membranes (12). Therefore, the loss of RAB18 does not affect normal BBSome targeting to the basal bodies but prevents its passage through the TZ for ciliary entry (12). Given that the absence of RAB23 results in a similar failure of BBSome entry into cilia, we investigated whether RAB23 facilitates BBSome passage through the TZ by activating RAB18. To address this, we analyzed the RAB18 KD strain *RAB18^miR^* and the corresponding rescue strains *RAB18^miR^; RAB18:HA:YFP-TG*, *RAB18^miR^; RAB18Q67L:HA:YFP-TG*, and *RAB18^miR^; RAB18S22N:HA:YFP-TG* (12). These strains express RAB18-HA-YFP in its wild-type, GTP-bound (Q67L), and GDP-bound (S22N) states, respectively, at levels comparable to the endogenous RAB18 in the parental CC-5325 cells (12). RAB18 KD did not alter the cellular levels of the BBSome or RAB23 (Fig. 3*A*), nor their localization to the IFT81-labeled basal bodies (Fig. 3*B* and *SI Appendix,* Fig. S6). However, the BBSome and RAB23 almost entirely failed to enter cilia of *RAB18^miR^*and *RAB18^miR^; RAB18S22N:HA:YFP-TG* cells, whereas it entered normally into cilia of *RAB18^miR^; RAB18:HA:YFP-TG* and *RAB18^miR^; RAB18Q67L:HA:YFP-TG* cells (Fig. 3*A*). This result supports the requirement for RAB18-GTP in facilitating the BBSome’s passage across the TZ for ciliary entry (12) and its simultaneous role in enabling RAB23-GDP passage across TZ for ciliary entry. (Fig. 2*E*). Notably, consistent with our previous study, the RAB18Q67L mutant promoted BBSome entry into cilia (Fig. 3*A*) (12). However, the BBSome failed to associate with anterograde IFT trains, instead accumulating at the proximal ciliary base in *RAB18^miR^; RAB18Q67L:HA:YFP-TG* cells (Fig. 3*B*) (12). In this strain, RAB23 entered cilia (Fig. 3*A*) but did not accumulate in the same proximal ciliary base as the BBSome (*SI Appendix,* Fig. S6), suggesting that RAB23-GDP may transition to its GTP-bound state to facilitate its release from the BBSome.

**Figure 3.**
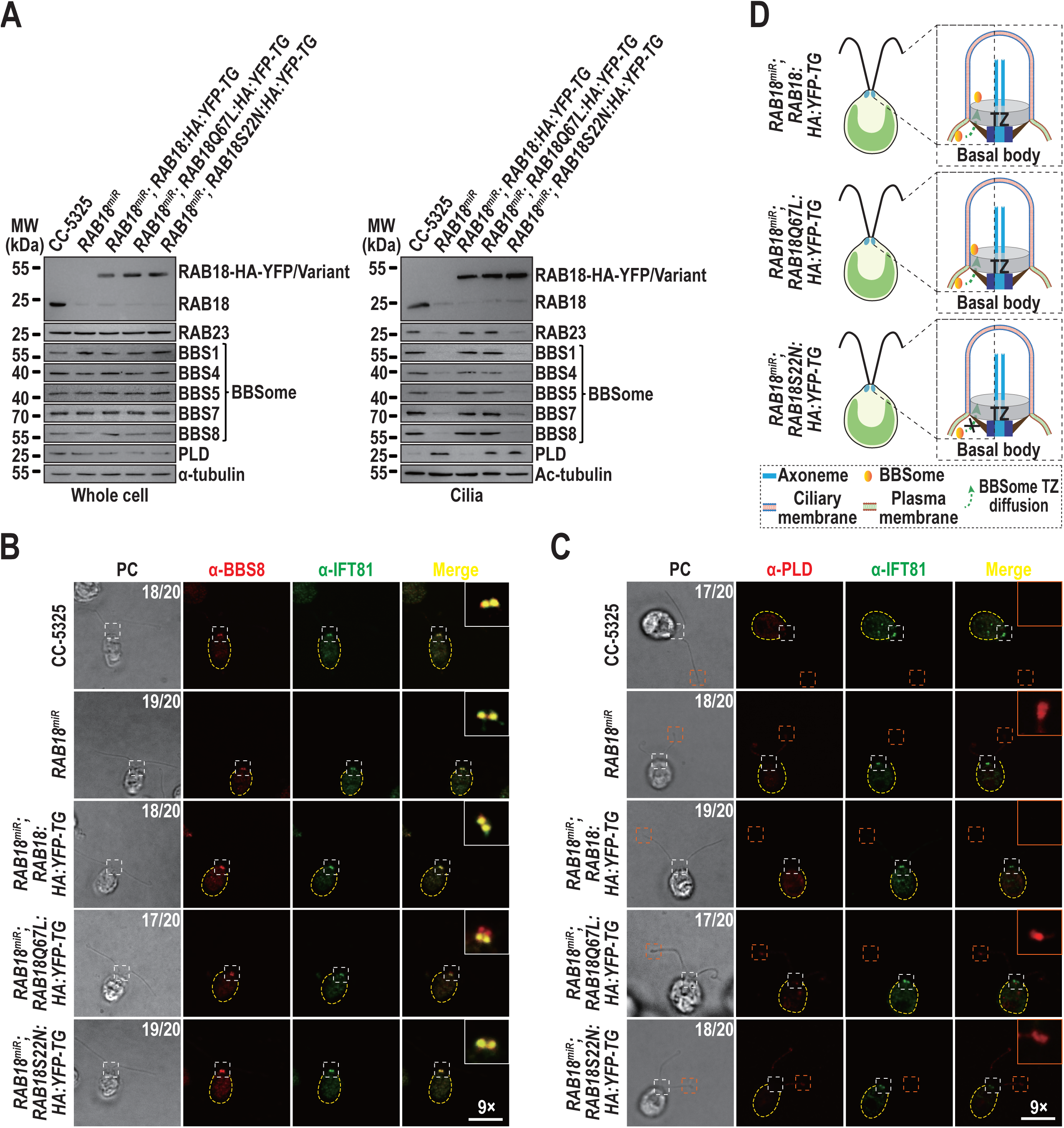
RAB23-GDP facilitates inward BBSome TZ passage by activating RAB18. (*A*). Immunoblots of whole cell samples and cilia from the indicated strains (top) probed with α-RAB18, α-RAB23, α-BBS1, α-BBS4, α-BBS5, α-BBS7, α-BBS8, and α-PLD. Alpha-tubulin and Ac-tubulin served as loading controls for whole cell samples and cilia, respectively. MW indicates molecular weight. (*B*). Immunofluorescence staining of the indicated cells (left) with α-BBS8 (red) and α-IFT81 (green). Cell counts (out of 20) represents cells positive for BBS8 and IFT81 at the basal bodes and negative or positive for BBS8 at the TZ. (*C*). Immunofluorescence staining of the indicated cells (left) with α-PLD (red) and α-IFT81 (green). Cell counts (out of 20) represents cells positive or negative for PLD at the ciliary tip. (*D*) Schematic representation of the mechanism by which RAB23 activates RAB18 to promote the inward TZ passage of the BBSome. For panels *B* and *C*, representative PC images are shown. Insets are 9 times magnifications of the basal body and TZ region indicated with a gray box. Scale bar: 10 µm.

Our previous studies demonstrated that ciliary membrane-anchored PLD undergoes TZ crossing for ciliary retrieval via the RABL2-ARL3 module-mediated outward BBSome TZ diffusion pathway (15, 16). In *RAB18^miR^* and *RAB18^miR^; RAB18S22N:HA:YFP-TG* cilia, the absence of the BBSome disrupted PLD-BBSome coupling at the ciliary tip, leading to PLD retention within cilia, particularly at the tip, where it became detectable by immunofluorescence staining (Fig. 3*A* and 3*C*) (15, 16). So did for *RAB18^miR^; RAB18Q67L:HA:YFP-TG* cilia as the “GTP-locked” RAB18Q67L mutant failed to hydrolyze GTP, preventing BBSome release from the proximal ciliary membrane, integration into anterograde IFT trains, and subsequent transport to the ciliary tip for PLD coupling (Fig. 3*A* and 3*C*) (12, 15, 16). In contrast, PLD dynamics remains normal in *RAB18^miR^; RAB18:HA:YFP-TG* cilia, where BBSome homeostasis is preserved (Fig. 3*A* and 3*C*). Since RAB18-GTP, by being anchored to the cell membrane, binds and recruits the BBSome to cross the TZ for ciliary entry (12), these findings suggest that RAB23-GDP activates RAB18-GDP at the basal bodies to promote inward BBSome TZ passage. Under physiological conditions, RAB18-GDP localizes to the basal bodies, where it converts to RAB18-GTP under RAB23-GDP mediation (Fig. 3*D*).

### RAB23-GDP functions as a GEF for RAB18

Functional studies indicate that RAB23-GDP facilitates RAB18 from its GDP-bound to GTP-bound state at the basal bodies, suggesting that RAB23 acts as a GEF for RAB18 (Fig. 3). To test this hypothesis, we performed immunoprecipitation assays on cell body extracts of the BBSome-deficient strains, including *RAB18^miR^; bbs8; RAB18:HA:YFP-TG*, *RAB18^miR^; bbs8; RAB18Q67L:HA:YFP-TG*, and *RAB18^miR^; bbs8; RAB18S22N:HA:YFP-TG*, with HA-YFP-expressing HR-YFP cell serving as a negative control (12, 36). In the presence of the nonhydrolytic GTP analog GTPγS, which locks both RAB23 and RAB18 in their GTP-bound states, neither RAB18-HA-YFP nor its variants recovered RAB23 (Fig. 4*A*). However, in the presence of GDP, which maintains both GTPases in their GDP-bound states, RAB18-HA-YFP and RAB18S22N-HA-YFP, but not RAB18Q67L-HA-YFP successfully recovered RAB23 (Fig. 4*A*). These results reveal that RAB23 interacts efficiently with RAB18 only when both are GDP-bound, and that this interaction occurs independently of the BBSome.

**Figure 4.**
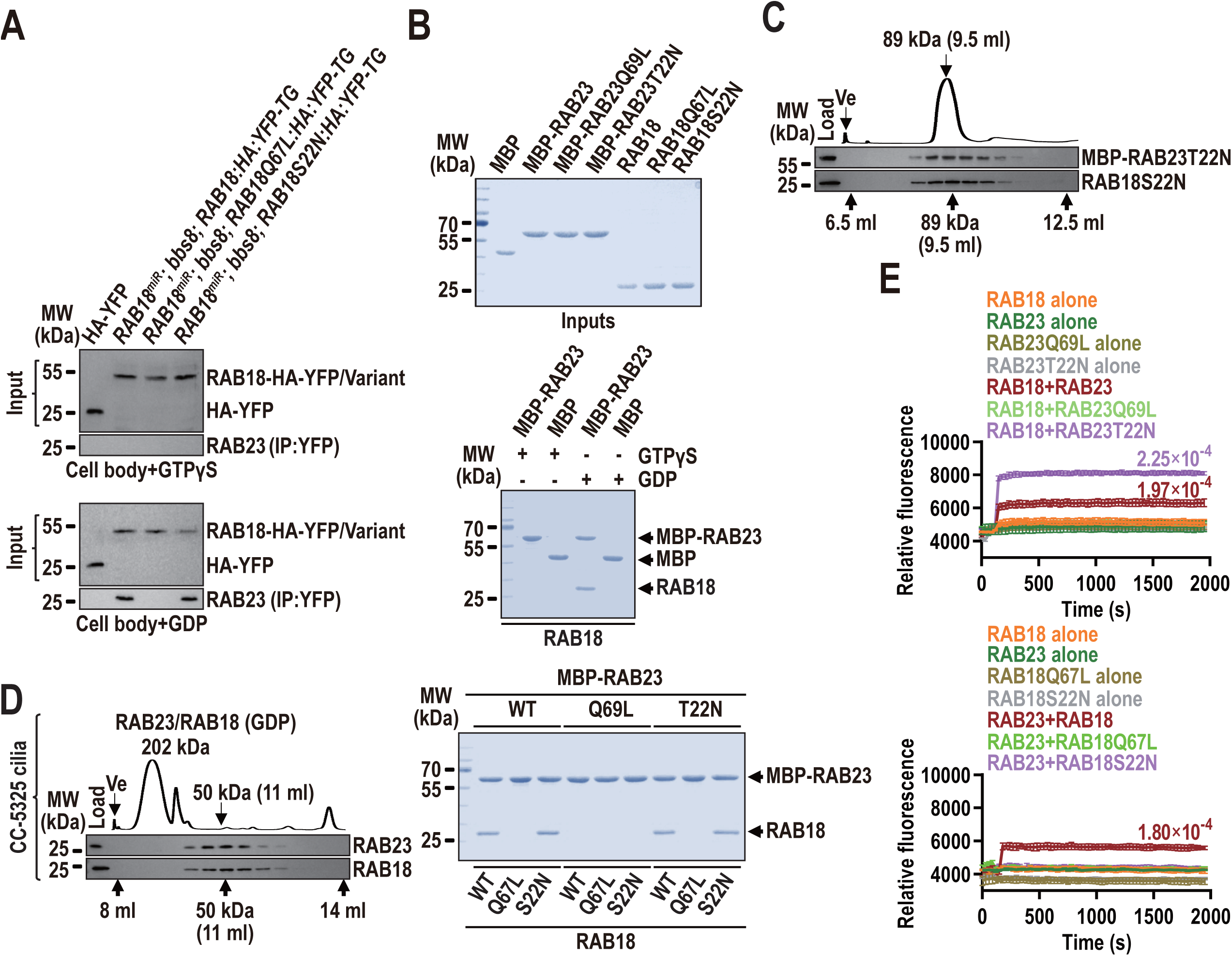
RAB23-GDP functions as a GEF for RAB18. (*A*). Immunoblots of α-YFP-captured proteins from cell body of the indicated strains (top) in the presence of GTPγS or GDP probed with α-RAB23. The input was adjusted with α-YFP by immunoblotting. (*B*). SDS-PAGE and Coomassie staining of MBP, MBP-RAB23, MBP-RAB23Q69L, MBP-RAB23T22N, RAB18, RAB18Q67L, and RAB18T22N (top). MBP alone and MBP-RAB23 were mixed with RAB18 in the presence of GTPγS or GDP (middle). Nine combinations of the above six proteins were mixed (bottom). Complexes recovered on amylose beads were resolved by SDS-PAGE and visualized by Coomassie staining. (*C-D*). The MBP-RAB23T22N/RAB18S22N complex recovered on amylose beads (*C*) or cell body samples of C-5325 in the presence of GDP (*D*) were fractionated by size exclusion chromatography (S200 sizing column) and probed with α-RAB23 and α-RAB18. Relative protein concentrations of the fractions between elution volumes of 6.5-12.5 ml (*C*) and 8-14 ml (*D*) were recorded as absorbance at 280 nm. The excluded volumes (Ve) and the elution volumes of the MBP-RAB23T22N/RAB18S22N complex (89 kDa) (*C*) and RAB23/RAB18 in the presence of GDP (50 kDa) (*D*) are indicated under the immunoblots. In panel *D*, 202 kDa refers to unknown *Chlamydomonas* ciliary protein complexes. (*E*). Mant-GTP association measurements to determine RAB23 GEF activity on RAB18. Fluorescence intensity of mant-GTP (0.75 µM) was plotted against recording time (seconds) for the following concentrations: 2 µM RAB18, RAB23Q69L, or RAB23T22N alone, or in the presence of 0.5 µM RAB23Q69L or RAB23T22N (top, GEF activity on RAB18); and 2 µM RAB23, RAB18Q67L, or RAB18S22N alone, or in the presence of 0.5 µM RAB18Q67L or RAB18S22N (bottom, GEF activity on RAB23). Data represent mean ± S.D. from three replicates. For panels *A*, *B*, *C*, *D*, MW indicates molecular weight.

To investigate whether RAB23-GDP directly interacts with RAB18-GDP, we prepared bacterially expressed maltose-binding protein (MBP), N-terminal 6×His-tag-removed RAB18, RAB18Q67L, and RAB18S22N, and N-terminal MBP-tagged RAB23, RAB23Q69L, and RAB23T22N (Fig. 4*B*). *In vitro* interaction assays excluded the interaction of MBP with RAB18 in either nucleotide state (Fig. 4*B*) and confirmed that interaction between the two GTPases occur only when both are in a GDP-bound state (Fig. 4*B*). Additionally, the assays revealed that bacterially expressed RAB18 and RAB23 predominantly exist in their GDP-bound states (Fig. 4*B*). Size exclusion chromatography of the pulldown eluate demonstrated that the binding of MBP-RAB23T22N and RAB18S22N occurs at a molecular ratio of 1:1, with a combined molecular weight of approximately 89 kDa (Fig. 4*C*). Similar result was obtained using flagellar extracts from CC-5325 cells in the presence of GDP (Fig. 4*D*). GEF assays further confirmed that bacterially expressed RAB23T22N and RAB23 promotes RAB18 to bind mant-GTP at a rate of 2.25×10^-4^ and 1.97×10^-4^ (µmol mant-GTP/µmol RAB18/sec), respectively, whereas RAB23Q69L failed to do so, establishing RAB23 as a GEF for RAB18 (Fig. 4*E*). In contrast, neither RAB18S22N nor RAB18Q67L promoted RAB23 to bind mant-GTP, ruling out RAB18 as a GEF for RAB23 (Fig. 4*E*). The conclusion was further supported by mant-GDP dissociation assays in the presence of the non-hydrolytic GTP analogue GppNHp, where RAB23S22N and RAB23 facilitated the release of GDP from RAB18 with a rate constant (*Kobs*) of 3.31×10^-3^/s and 2.64×10^-3^/s, respectively (*SI Appendix,* Fig. S7*A*). In summary, we propose that RAB23-GDP transiently binds RAB18-GDP to activate it as a GEF at the basal bodies, enabling its transition to RAB18-GTP. Importantly, this activation is unidirectional, as RAB18 does not act as a GEF for RAB23.

### RAB23-GDP binds to RAB18-GTP/BBSome as a BBSome cargo for inward TZ passage

The failure of RAB23-GDP to enter cilia in the absence of the BBSome suggests that it enters cilia through the RAB18-mediated BBSome ciliary entry process (Fig. 2*E*). To testify this hypothesis, we performed immunoprecipitation assays on cell body extracts from *RAB23^miR^; RAB23:HA:YFP-TG* cells. In the absence of GDP or GTPγS, RAB23-HA-YFP immunoprecipitated both RAB18 and the BBSome (Fig. 5*A*). However, when GTPγS was present, which renders RAB23 and RAB18 in their GTP-bound states, RAB23-HA-YFP failed to recover either RAB18 or the BBSome (Fig. 5*A*). In contrast, in the presence of GDP, which retains both GTPases in their GDP-bound states, RAB23-HA-YFP recovered both RAB18 and the BBSome (Fig. 5*A*). These findings suggest that RAB23-GTP does not interact with the BBSome of either RAB18-GTP-bound or RAB18-free. Instead, RAB23-GDP binds simultaneously to RAB18-GDP and the BBSome, free from RAB18-GTP binding. Supporting this notion, RAB18Q67L-HA-YFP immunoprecipitated both the BBSome and RAB23 only when RAB23 was rendered GDP-bound with the addition of GDP (Fig. 5*B*). Conversely, RAB18S22N-HA-YFP recovered RAB23 preloaded with GDP but not GTPγS and failed to immunoprecipitate the BBSome under any nucleotide conditions (Fig. 5*B*). Further confirmation came from co-sediment assays performed on cell body extracts. The RAB18Q67L mutant co-sedimented with the BBSome and the endogenous RAB23 only when GDP was preloaded (Fig. 5*C*). In contrast, the RAB18S22N mutant did not co-sedimented with the BBSome (Fig. 5*C*). As reflected by the *E. coli*-based *in vitro* protein interaction assays, BBS2 was identified as the BBSome subunit efficiently captured by RAB23T22N, confirming that RAB23-GDP binds the BBSome through direct interaction with BBS2 (Fig. 5*D*). While single-molecule visualization of RAB23 crossing the TZ for ciliary entry is currently not feasible (Fig. 1*D*), these data suggest that RAB23-GDP interacts with RAB18-GTP/BBSome as a BBSome cargo for inward TZ passage. This implies that RAB23-GDP likely moves across the TZ for ciliary entry via the inward RAB18-GTP/BBSome TZ passage pathway (12).

**Figure 5.**
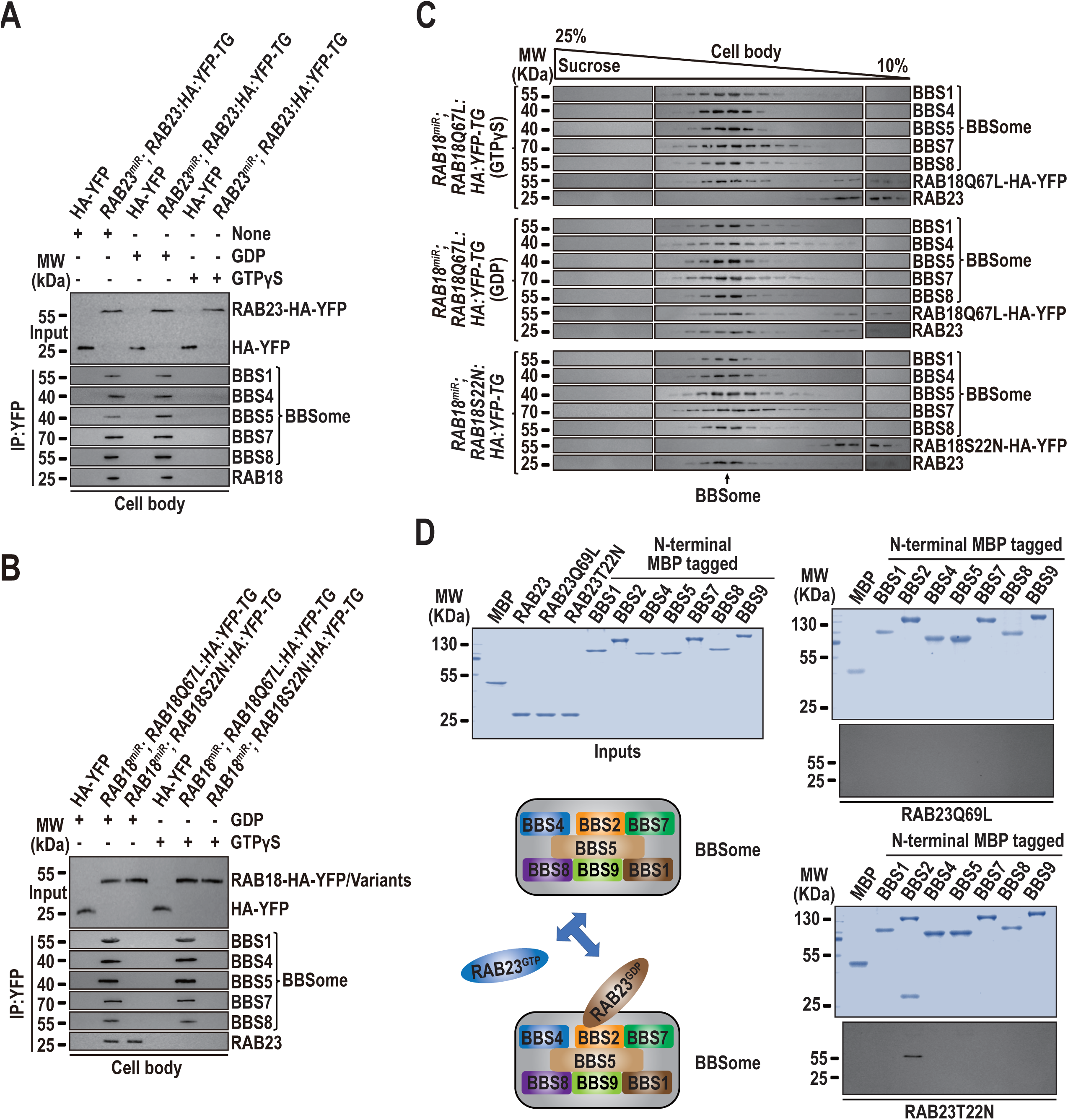
RAB23-GDP binds to RAB18-GDP/BBSome as a BBSome cargo for inward TZ passage. (*A*). Immunoblots of α-YFP-captured proteins from cell body of the indicated strains (top) in the presence of GDP, GTPγS, or no nucleotide probed with α-BBS1, α-BBS4, α-BBS5, α-BBS7, α-BBS8, and α-RAB18. (*B*). Immunoblots of α-YFP-captured proteins from cell body of the indicated strains (top) in the presence of GDP or GTPγS probed with α-BBS1, α-BBS4, α-BBS5, α-BBS7, α-BBS8, and α-RAB23. (*C*). Immunoblots of sucrose density gradient fractions from cell body samples of the indicated strains (left) in the presence of GTPγS, GDP or no nucleotide probed with α-BBS1, α-BBS4, α-BBS5, α-BBS7, α-BBS8, α-RAB18, and α-RAB23. (*D*). Bacterially expressed MBP, MBP-BBS1, MBP-BBS2, MBP-BBS4, MBP-BBS5, MBP-BBS7, MBP-BBS8, and MBP-BBS9 (top left) were mixed with RAB23Q69L or RAB23T22N (top left). Complexes recovered on amylose beads were resolved by SDS-PAGE, followed by Coomassie staining and immunoblotting with α-RAB23 (right). A schematic representation of the direct interaction between RAB23-GDP with the BBS2 subunit of the BBSome is shown (bottom left). For panels *A* and *B*, input levels were normalized using α-YFP immunoblotting. For all panels, MW denotes molecular weight.

### Mechanistic characterization of RAB23 mutations on impeding inward BBSome TZ passage

Carpenter syndrome (CS) patients carrying the RAB23C85R mutation (equivalent to the A86R mutation in *C. reinhardtii*) present with craniosynostosis, obesity, polydactyly, and soft-tissue syndactyly, with some symptoms overlapped with those of BBS (37–39). These patients exhibit defects in Shh signaling to varying degrees (38). In our *Chlamydomonas* model, the BBSome as an integrated complex fails to pass the TZ for ciliary entry when RAB23 is absent, as RAB23 is required to activate RAB18 at the basal bodies. If this mechanism is conserved in humans, the loss of RAB23 would render the BBSome unavailable to couple with Shh signaling GPCRs such as Smo, GPR161, and Patch1 at the ciliary tip, ultimately impairing Hh signaling pathways (40, 41). To investigate how the RAB23A86R mutation impacts BBSome ciliary entry, we expressed RAB23A86R-HA-YFP at levels comparable to endogenous RAB23 in the strain *RAB23^miR^*, generating the strain *RAB23^miR^; RAB23A86R:HA:YFP-TG* (Fig. 6*A*). The predominant replacement of endogenous RAB23 with RAB23A86R-HA-YFP did not affect cellular abundance of the BBSome (Fig. 6*A*) and the assembly of the BBSome in the cytoplasm (*SI Appendix,* Fig. S8*A*). It also did not disrupt BBSome targeting to the basal bodies (Fig. 6*B*). However, the mutation severely impaired BBSome entry into cilia (Fig. 6*A*). In addition, the A86R mutation did not affect RAB23 targeting to the basal bodies as labeled with IFT81 (Fig. 6*C*). However, RAB23A86R-HA-YFP failed to immunoprecipitate the BBSome or RAB18, even in the presence of GDP (*SI Appendix,* Fig. S8*B*). This observation was corroborated by *E. coli*-based protein-protein interaction assays, which demonstrated that the A86R mutation abolishes the GDP-dependent interaction between RAB23 and RAB18 (Fig. 6*D*). This eliminated RAB23’s GEF activity on RAB18 (Fig. 6*E* and *SI Appendix,* Fig. S7*B*). In summary, RAB23A86R targets the basal bodies similarly to RAB23 but, unlike RAB23, fails to interact with RAB18 and cannot catalyze the conversion of RAB18-GDP to RAB18-GTP at the basal bodies. It also loses its ability to interact with the BBSome as a BBSome cargo, thereby failing to enter cilia (*SI Appendix,* Fig. S8*A* and *B*).

**Figure 6.**
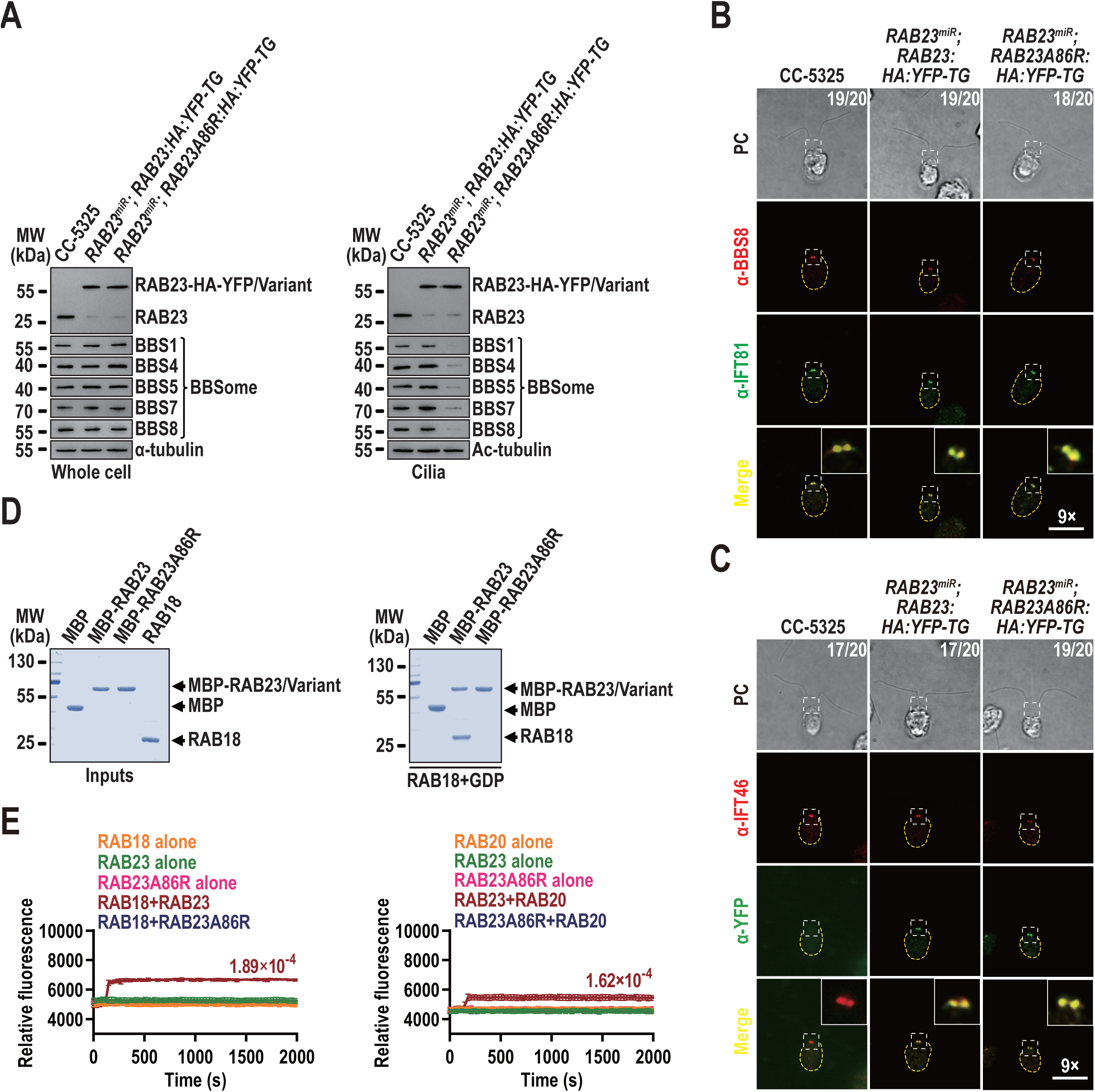
Mechanistic characterization of RAB23 mutations on impeding inward BBSome TZ passage. (*A*). Immunoblots of whole cell samples and cilia from the indicated strains (top) probed with α-RAB23, α-BBS1, α-BBS4, α-BBS5, α-BBS7, and α-BBS8. Alpha-tubulin and Ac-tubulin were used to normalize the loading for whole cell samples and cilia, respectively. MW indicates molecular weight. (*B*). Immunofluorescence staining of the indicated cells (top) with α-BBS8 (red) and α-IFT81 (green). Cell counts out of 20 are listed, indicating the number of cells positive for IFT81 and BBS8 at the basal bodies. (*C*). Immunofluorescence staining of the indicated cells (top) with α-IFT46 (red) and α-YFP (green). Cell counts out of 20 are listed, indicating the number of cells positive for IFT46 and negative or positive for RAB23-HA-YFP and RAB23A86R-HA-YFP at the basal bodies. (*D*). Bacterially expressed MBP, MBP-RAB23, MBP-RAB23A86R, and RAB18 were purified, resolved by SDS-PAGE, and visualized by Coomassie staining (left). RAB18 was mixed with MBP, MBP-RAB23, and MBP-RAB23D76G. Complexes recovered on amylose beads were resolved by SDS-PAGE and visualized by Coomassie staining (right). (*E*). Mant-GTP association assays were conducted to assess RAB23A86R GEF activity on RAB18. Fluorescence intensity of mant-GTP (0.75 µM) was plotted against recording time (seconds) for the following concentrations: 2 µM RAB18, RAB23, or RAB23A86R alone or in the presence of 0.5 µM RAB23 or RAB23A86R (left, GEF activity on RAB18); and 2 µM RAB23, RAB18, or RAB23A86R alone, or in the presence of 0.5 µM RAB18 (right, GEF activity on RAB23). Data are shown as mean ± S.D. from three replicates. For panels *B* and *C*, representative PC images are shown. Insets are 9 times magnifications of the basal body and the TZ region indicated with a gray box. Scale bar: 10 µm.

## Discussion

In this study, *C. reinhardtii* was employed as a model organism to demonstrate that RAB23, only in a GDP-bound state, targets the basal bodies. Upon reaching the basal bodies, RAB23-GDP functions as a GEF for RAB18, converting RAB18-GDP into RAB18-GTP. The active RAB18-GTP then anchors to the plasma membrane and binds and recruits the BBSome as its effector to diffuse through the TZ via lateral membrane transportation into cilia (12). RAB23-GDP crosses the TZ to enter cilia, but it does so by binding to the RAB18-GTP-bound BBSome as a cargo (Fig. 7). Our findings offer critical insights into the interplay between the small GTPases RAB23 and RAB18 and their roles in guiding the BBSome across the TZ for ciliary entry. Additionally, they address a key gap in understanding how this RAB23-RAB18 module facilitates the export of BBSome cargoes, such as PLD, from cilia in *C. reinhardtii*.

**Figure 7.**
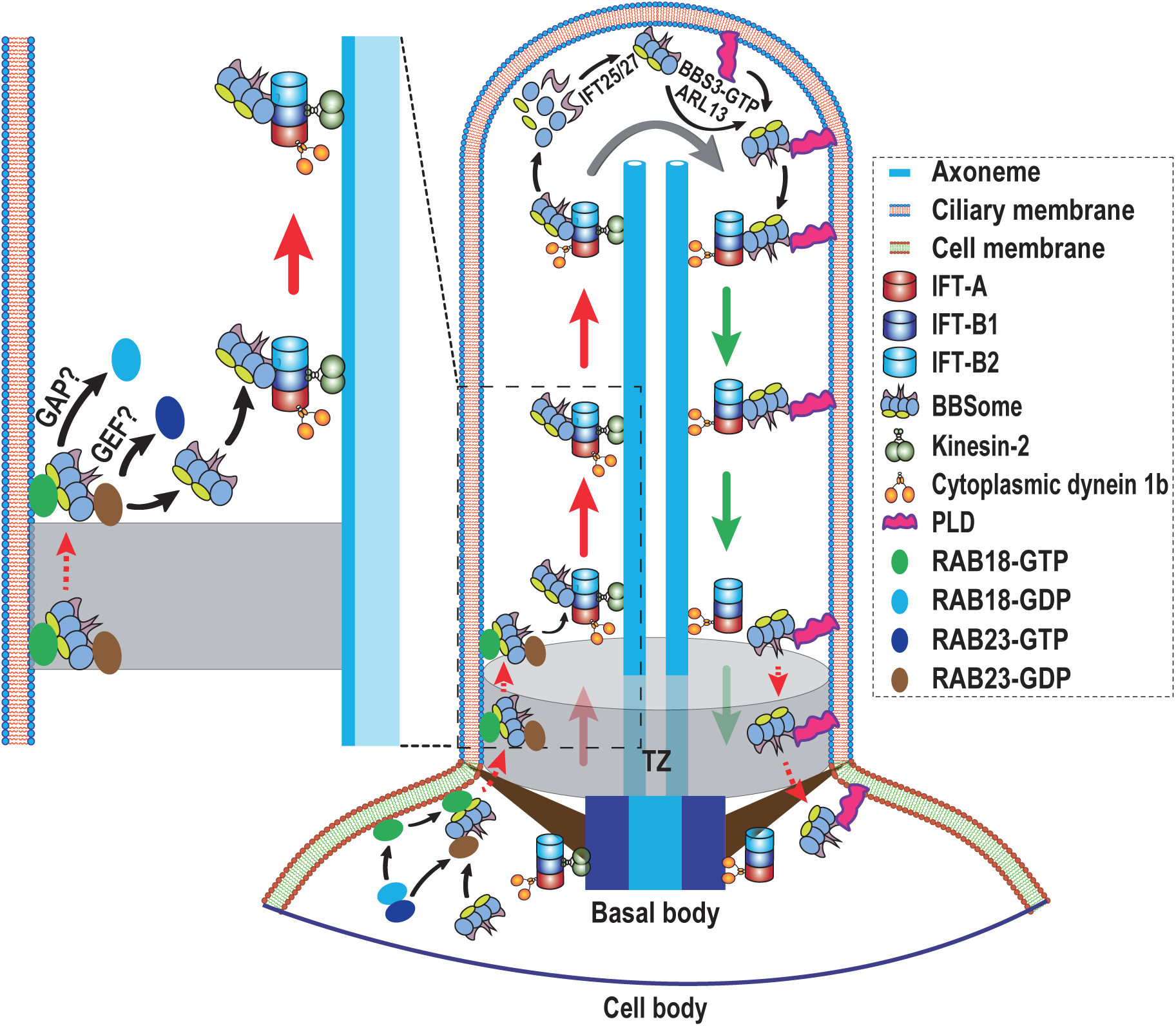
Hypothetical model for how *Chlamydomonas* RAB23 promotes BBSome diffusion through the TZ for ciliary entry through activating RAB18. In a basal body region just below the TZ, RAB23 in its GDP-bound form acts as a GEF for RAB18, converting RAB18-GDP to its active GTP bound state. Anchored to the plasma membrane, RAB18-GTP binds its BBSome effector independently of IFT association. Subsequently, RAB18-GTP facilitate the lateral diffusion of the BBSome, allowing its entry into cilia. During this process, RAB23-GDP is transported into cilia by coupling with the RAB18-GTP/BBSome comple as a BBSome cargo. This transport is mediated through a direct interaction between RAB23-GDP and the BBSome subunit BBS2. Upon reaching the proximal ciliary base right above the TZ, an unidentified GTPase-activating protein (GAP) hydrolyzes RAB18-GTP, leading to the release of the BBSome from the ciliary membrane before its integration into anterograde IFT trains. At this stage, an unknown GEF is presumed to convert RAB23-GDP into its active GTP-bound form, facilitating its release from the BBSome. Upon reaching the ciliary tip, the BBSome dissociates with anterograde IFT trains for residing in the ciliary tip matrix and remodels with the aid of the IFT-B1-shed IFT25/27 (13, 14). The remodeled BBSome then couples with its cargo PLD via the BBS3-ARL13 module (23, 24). This interaction enables the PLD-laden BBSome to execute a U turn at the ciliary tip and undergoes retrograde transport toward the ciliary base via retrograde IFT (23). Upon reaching the proximal ciliary base right above the TZ, the PLD-laden BBSome separates from retrograde IFT trains and diffuses through the TZ for ciliary retrieval, a process mediated by the RABL2-ARL3 module (15, 16).

### RAB23 activates RAB18 at the basal bodies for promoting BBSome crossing the TZ for ciliary entry

At physiological conditions, *Chlamydomonas* RAB18 rapidly transitions from a GDP-bound to a GTP-bound state at the basal bodies (12). This conversion enables RAB18-GTP to anchor to the plasma membrane, facilitating its diffusion through the TZ and entry into cilia via lateral membrane transport between the plasma and ciliary membranes (12). During this process, RAB18 interacts with the BBSome as its effector, recruiting it to pass through the TZ for ciliary entry (12). This function positions RAB18 as a key regulator of BBSome-dependent transmembrane and membrane anchoring signaling protein (such as PLD) homeostasis in cilia (12). However, the mechanism underlying RAB18 nucleotide conversion at the basal bodies has remained unclear. Our study addresses this gap by demonstrating that RAB23-GDP functions as a RAB18 GEF at the basal bodies (Fig. 4). According to our data, GDP-bound RAB23 localizes to a region just below the TZ, where it activates RAB18-GDP, converting it into its GTP-bound form (Fig. 7). While it remains uncertain whether this mechanism is conserved in mammalian and human Rab23, our findings suggest that in *C. reinhardtii*, RAB23 maintains BBSome homeostasis by regulating its availability for TZ-mediated ciliary entry via RAB18. In this way, it contributes to maintaining the homeostasis of ciliary signaling proteins in a BBSome-dependent manner.

### RAB23 likely undergoes nucleotide exchange in the proximal ciliary base

RAB23-GDP, which binds the BBSome via a direct interaction with the BBS2 subunit, diffuses into cilia via the RAB18-GTP/BBSome pathway (Fig. 3 and 5). Unlike IFT proteins that are easily visualized at the basal bodies and shown to undergo bidirectional IFT, RAB23 appears enriched at the basal bodies but does not participate in IFT (Fig. 1*A* and *C* and *SI Appendix,* Fig. S6). This observation indicates that, once reaching the proximal ciliary based localized right above the TZ, RAB23 must dissociate from the BBSome before its integration into anterograde IFT trains. Therefore, a nucleotide exchange from GDP to GTP must occur to RAB23, likely catalyzed by a RAB23-specific GTPase activating protein (GAP) (Fig. 7). However, the identity of this GAP remains to be determined. Our previous studies and others have shown that several GTPases exhibit multiple roles, functioning both inside and outside cilia (9, 11, 24). For example, ARL6/BBS3 binds to IFT22 in the cytoplasm to form an IFT22/BBS3 heterodimer in a nucleotide-independent manner in *C. reinhardtii* (11). It interacts with the BBSome and facilitates BBSome recruitment to the basal bodies when both are GTP-bound (11). A similar functional behavior of and the BBSome in this aspect was also observed in mammalian cells (9). Once inside cilia, BBS3 acts as a GEF for ARL13 at the ciliary tip, activating ARL13 to recruit the BBSome as its effector to the ciliary tip membrane for cargo coupling (23, 24). Our data has shown that RAB23 is not involved in IFT and ciliation (Fig. 1*A* and *SI Appendix,* Fig. S2*B*-*D*). However, it remains uncertain if RAB23, like ARL13, mediates ciliary signaling within cilia (24). Further investigation is needed to clarify its potential function.

### Implications for the molecular basis of CS caused by RAB23 mutation in humans

As a ciliary GTPase, Rab23 unlikely regulates IFT, as its depletion or loss does not affect cilia formation or length in various cultured mammalian cells lines, zebrafish (Kuppfers vesicle), trypanosomes, and mice (nodal cilia) (42, 43) as well as *C. reinhardtii* (Fig. 1*A* and *SI Appendix,* Fig. S3). Instead, Rab23 functions as an inhibitor of cilium-based Shh signaling in mouse (27, 28, 30, 44), and mutations in RAB23 result in CS in humans (37–39). In the most recent study, Rab23 deficiency was found to cause abnormality of primary cilia in mouse central neurons, eventually connecting CS to a ciliopathy (45, 46). In *C. reinhardtii*, the ciliary membrane-anchored signaling protein PLD associates with the BBSome at the ciliary tip and diffuses through the TZ for ciliary retrieval via the RABL2-ARL3 module-mediated outward BBSome TZ passage pathway (15, 16). The RAB23-RAB18 module regulates BBSome availability in cilia for PLD coupling and subsequently removal from cilia, as they are essential for promoting inward BBSome TZ passage and ciliary entry (12). If this mechanism applies to humans, RAB23 dysfunction could impair ciliary Shh singling by preventing BBSome entry into cilia, potentially causing CS (37–39). Supporting this, the *Chlamydomonas* RAB23 A86R mutation, which is equivalent to the human CS-causing RAB23 C85R mutation, impairs RAB23 GEF activity on RAB18 (Fig. 6 and *SI Appendix,* Fig. S7*B* and S8) (37, 38). Since defects in BBSome basal body-cilium cycling also lead to Bardet-Biedle syndrome (BBS) in humans (20, 21, 47), this impairment may explain the overlap of CS symptoms with BBS-like features, such as obesity and polydactyly (37, 38). Further studies are needed to elucidate the specific effects of RAB23 mutations on BBSome ciliary homeostasis and their role in human CS and BBS.

## Materials and Methods

### Antibodies, Chlamydomonas strains, and culture conditions

All antibodies used in this study are detailed in *SI Appendix,* Table S1. A rabbit-raised polyclonal antibody against RAB23 was custom-produced by Beijing Protein Innovation LLC (Beijing, China). The *Chlamydomonas* strains employed in this study are listed in *SI Appendix,* Table S2. Strains CC-5325 and *rab23-544* (LMJ.RY0402.185544) were purchased from the *Chlamydomonas* Library Project (CLiP, https://www.chlamylibrary.org/allMutants) (48). The following previously reported were also used: *bbs8* (49), *bbs8; BBS8:YFP-TG* (16), HA-YFP (50), *RAB18^miR^*, *RAB18^miR^; RAB18:HA:YFP-TG*, *RAB18^miR^; RAB18Q67L:HA:YFP-TG*, *RAB18^miR^; RAB18S22N:HA:YFP-TG, RAB18^miR^; bbs8; RAB18:HA:YFP-TG*, *RAB18^miR^; bbs8; RAB18Q67L:HA:YFP-TG*, *RAB18^miR^; bbs8; RAB18S22N:HA:YFP-TG* (12) and *ift46-1; IFT46:YFP-TG* (51). *Chlamydomonas* cells were grown at room temperature in Tris acetic acid phosphate (TAP) medium or minimal 1 (M1) medium under continuous light with constant aeration. Antibiotics were supplemented as required for specific strains: 20 µg/ml paromomycin (Sigma-Aldrich), 5 µg/ml bleomycin (Invitrogen), both antibiotics combined (10 µg/ml paromomycin and 5 µg/ml bleomycin), or a combination of three antibiotics (10 µg/ml paromomycin, 5 µg/ml bleomycin, and 10 µg/ml hydromycin).

A variety of experimental protocols were employed in this study, with most briefly outlined in the main text to enhance clarity and comprehension. Detailed descriptions of each protocol can be found in *SI Appendix*.

### Statistics

Statistical analyses were conducted using GraphPad Prism 8.0 (GraphPad Software). Data are presented as mean ± S.D., derived from three independently experiments, where “n” represents the sample size. A two-tailed *t*-test was used to compare protein expression levels and ciliary length. While data distribution was assumed to be normal, this assumption was not formally tested. “n.s.” indicates non-significant differences.

### Data, Materials, and Software Availability

All study data are included in the article and/or supporting information. All materials including plasmids, strains, and reagents are available upon request.

## Acknowledgements

Research reported in this publication was supported by the National Natural Science Foundation of China (32470741 to Z.-C.F.). We thank the stuff from the Core Facilities, the Fourth Affiliated Hospital of School of Medicine, and International School of Medicine, International Institutes of Medicine, Zhejiang University, for their technical support. The founder has no role in study design, data collection and analysis, decision to publish, or preparation of the manuscript.

## Author contributions

Z.-C.F. designed research; S.-N.S., W.-J.S. and Y.W. performed research; S.-N.S. and Z.-C.F. analyzed data; and Z.-C.F. wrote the paper.

