## Supplementary data for "RAB23 Modulates Signaling Protein Ciliary Homeostasis through Promoting RAB18-mediated Inward BBSome Transition Zone Passage"

### Figure S1

A

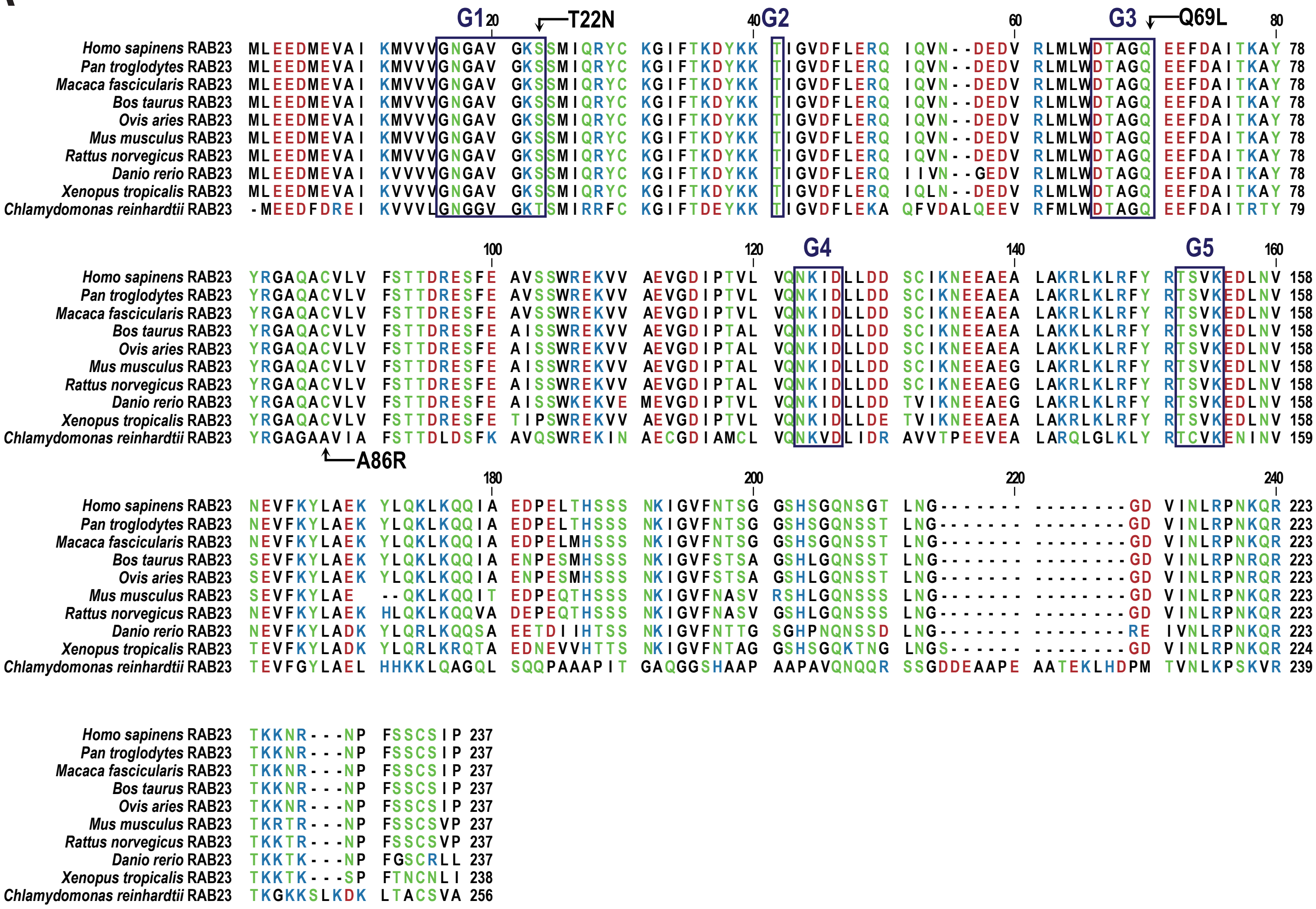


B

| Homology (vs. *C. reinhardtii* RAB23) | |
|---|---|
| *H. sapinens* | 50.58% |
| *P. troglodytes* | 50.58% |
| *M. fascicularis* | 50.58% |
| *B. taurus* | 49.42% |
| *O. aries* | 49.42% |
| *M. musculus* | 50.19% |
| *R. norvegicus* | 49.42% |
| *D. rerio* | 49.03% |
| *X. tropicalis* | 49.81% |

C

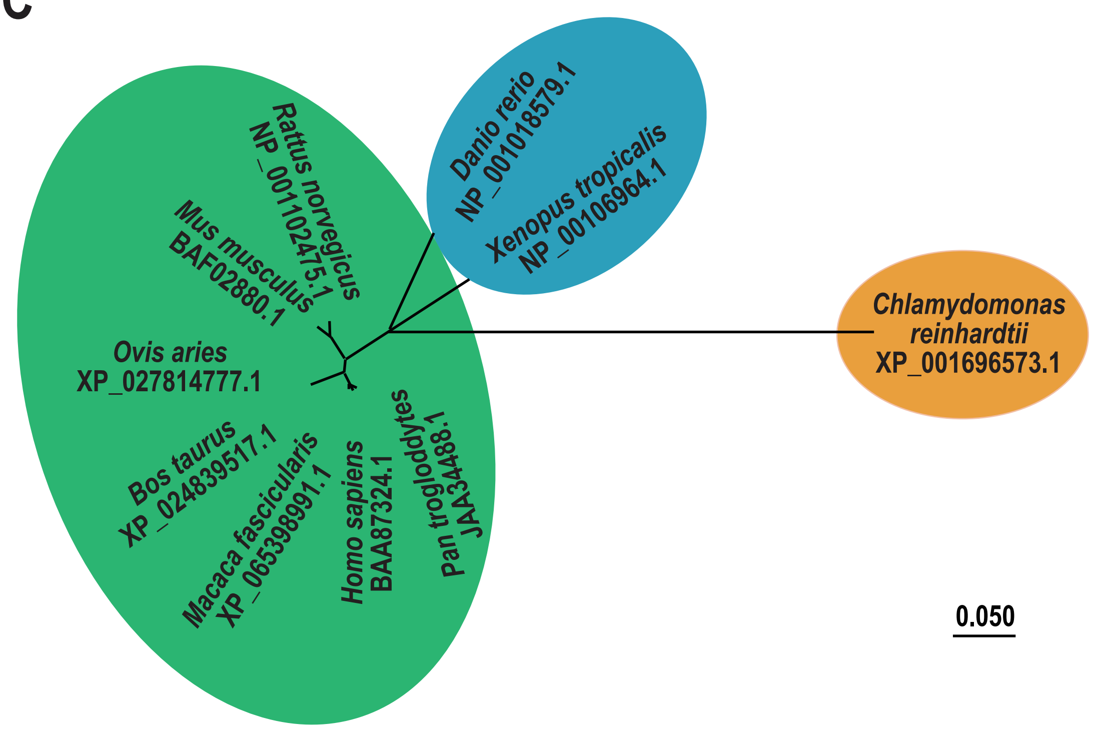

### Figure S2

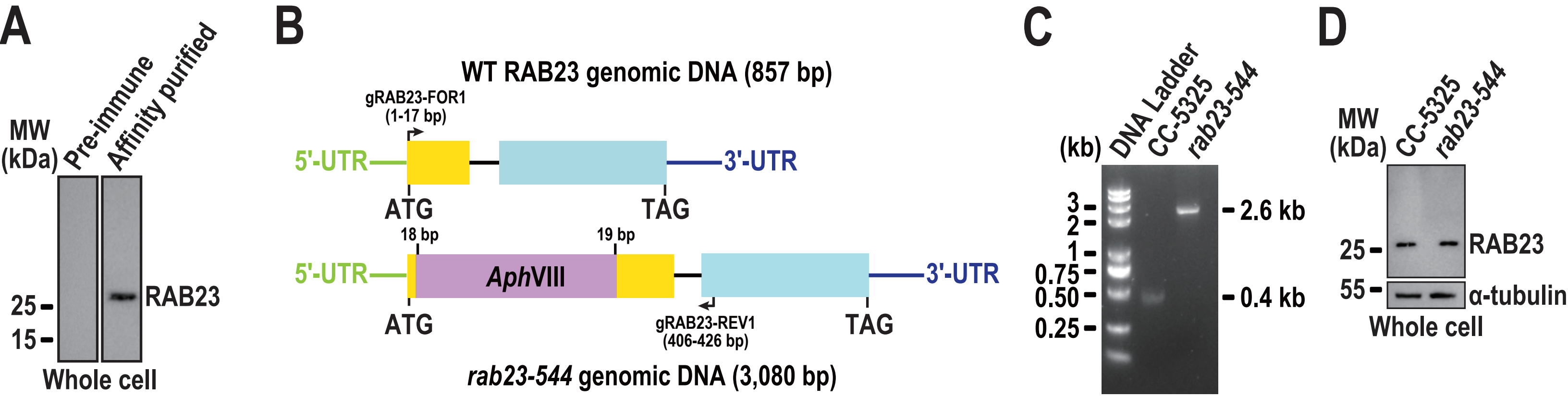
A
MW (kDa)
Pre-immune
Affinity purified
25
15
RAB23
Whole cell
B
WT RAB23 genomic DNA (857 bp)
gRAB23-FOR1 (1-17 bp)
5'-UTR
3'-UTR
ATG
TAG
18 bp
19 bp
AphVIII
gRAB23-REV1 (406-426 bp)
rab23-544 genomic DNA (3,080 bp)
C
(kb)
DNA Ladder
CC-5325
rab23-544
3
2
1
0.75
0.50
0.25
2.6 kb
0.4 kb
D
MW (kDa)
CC-5325
rab23-544
25
55
RAB23
α-tubulin
Whole cell

### Figure S3

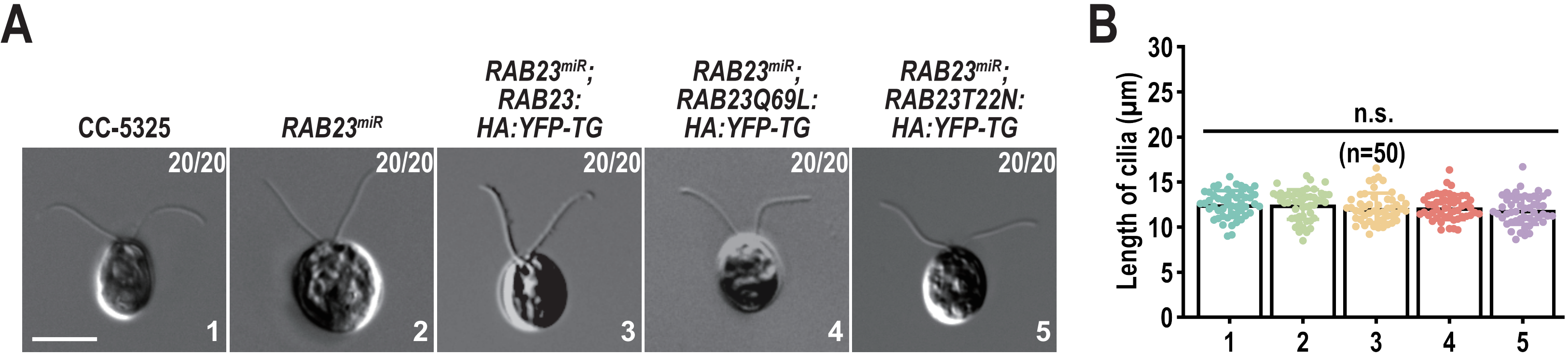
A
CC-5325
RAB23miR
RAB23miR; RAB23: HA:YFP-TG
RAB23miR; RAB23Q69L: HA:YFP-TG
RAB23miR; RAB23T22N: HA:YFP-TG
20/20
1
2
3
4
5
B
Length of cilia (μm)
n.s.
(n=50)
0
5
10
15
20
25
30

### Figure S4

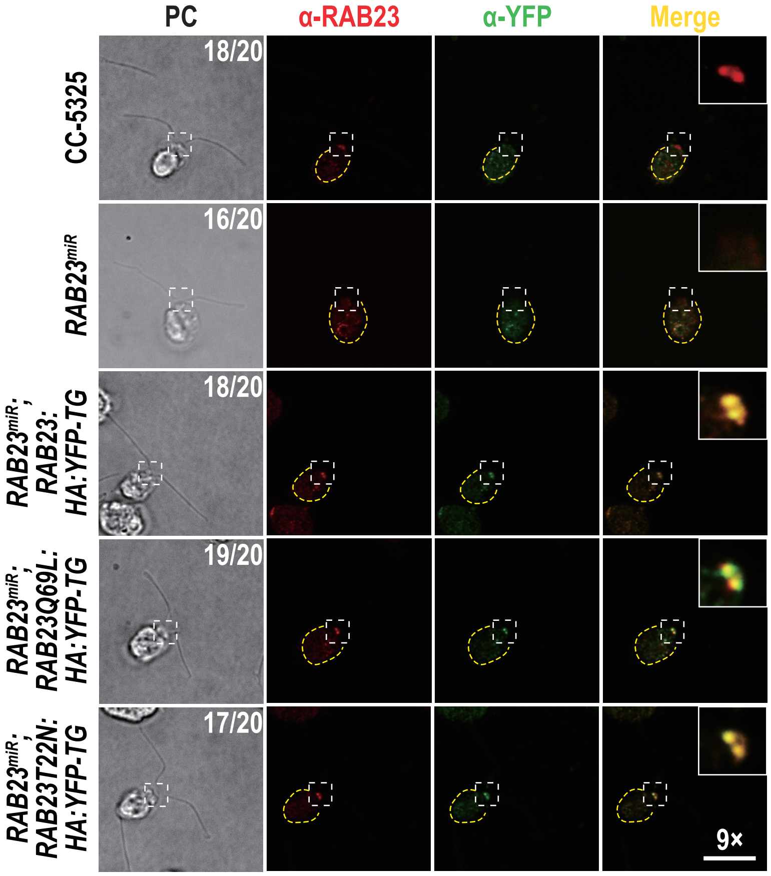
PC
α-RAB23
α-YFP
Merge
CC-5325
RAB23miR
RAB23miR; RAB23: HA:YFP-TG
RAB23miR; RAB23Q69L: HA:YFP-TG
RAB23miR; RAB23T22N: HA:YFP-TG
18/20
16/20
18/20
19/20
17/20
9×

### Figure S5

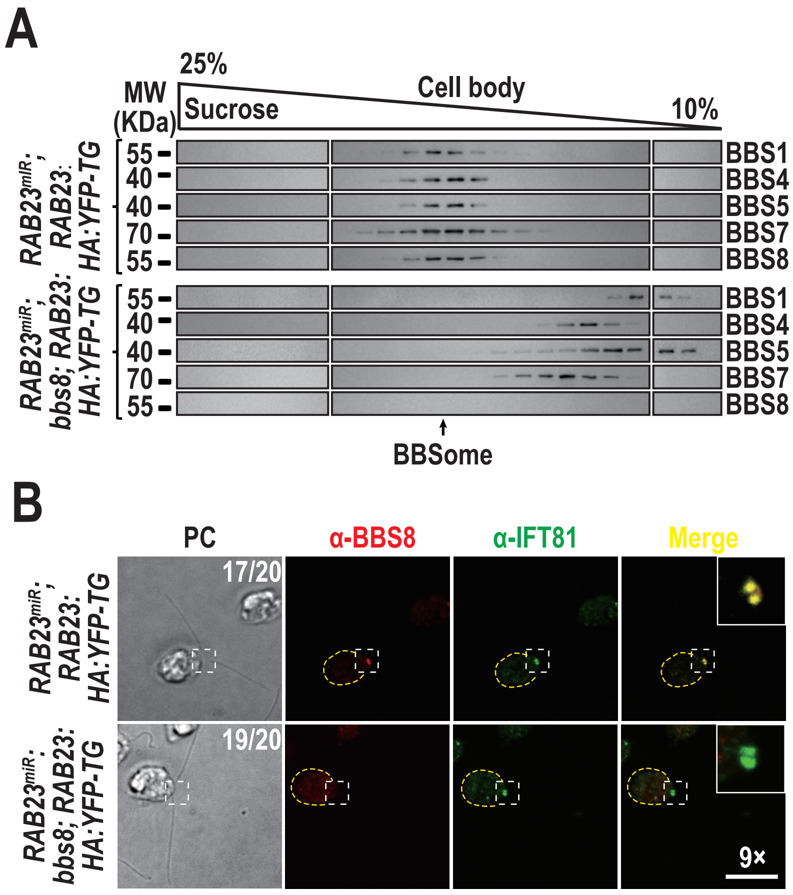
A
25%
Cell body
MW
(KDa)
Sucrose
10%
RAB23miR; RAB23: HA:YFP-TG
55
40
40
70
55
BBS1
BBS4
BBS5
BBS7
BBS8
RAB23miR; bbs8; RAB23: HA:YFP-TG
55
40
40
70
55
BBS1
BBS4
BBS5
BBS7
BBS8
BBSome
B
PC
α-BBS8
α-IFT81
Merge
RAB23miR; RAB23: HA:YFP-TG
17/20
RAB23miR; bbs8; RAB23: HA:YFP-TG
19/20
9×

### Figure S6

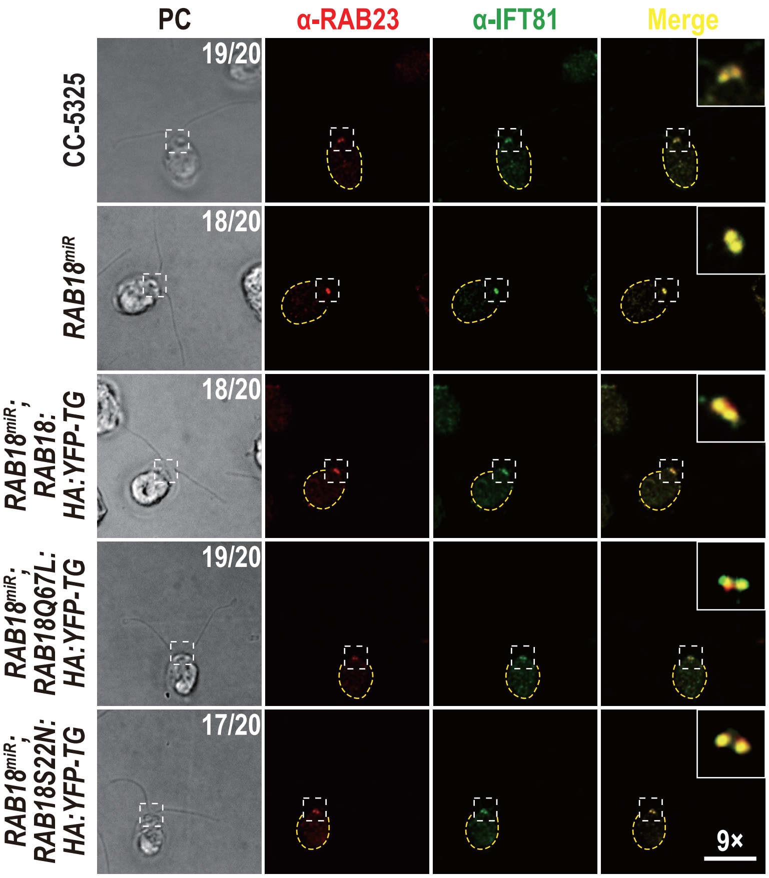

PC
α-RAB23
α-IFT81
Merge
CC-5325
RAB18miR
RAB18miR;
RAB18:
HA:YFP-TG
RAB18miR;
RAB18Q67L:
HA:YFP-TG
RAB18miR;
RAB18S22N:
HA:YFP-TG
19/20
18/20
18/20
19/20
17/20
9×

### Figure S7

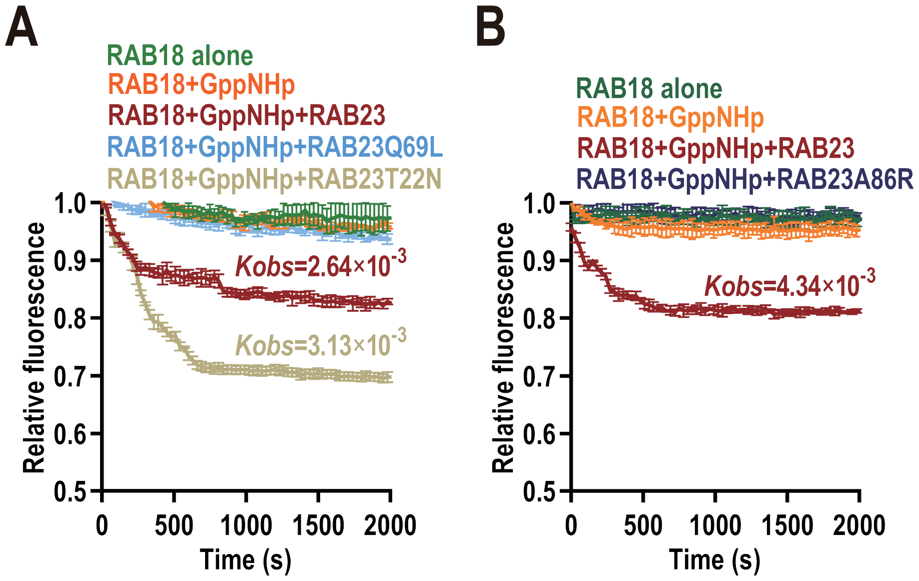
A
RAB18 alone
RAB18+GppNHp
RAB18+GppNHp+RAB23
RAB18+GppNHp+RAB23Q69L
RAB18+GppNHp+RAB23T22N
Kobs=2.64×10-3
Kobs=3.13×10-3
Relative fluorescence
1.0
0.9
0.8
0.7
0.6
0.5
0
500
1000
1500
2000
Time (s)
B
RAB18 alone
RAB18+GppNHp
RAB18+GppNHp+RAB23
RAB18+GppNHp+RAB23A86R
Kobs=4.34×10-3
Relative fluorescence
Time (s)

### Figure S8

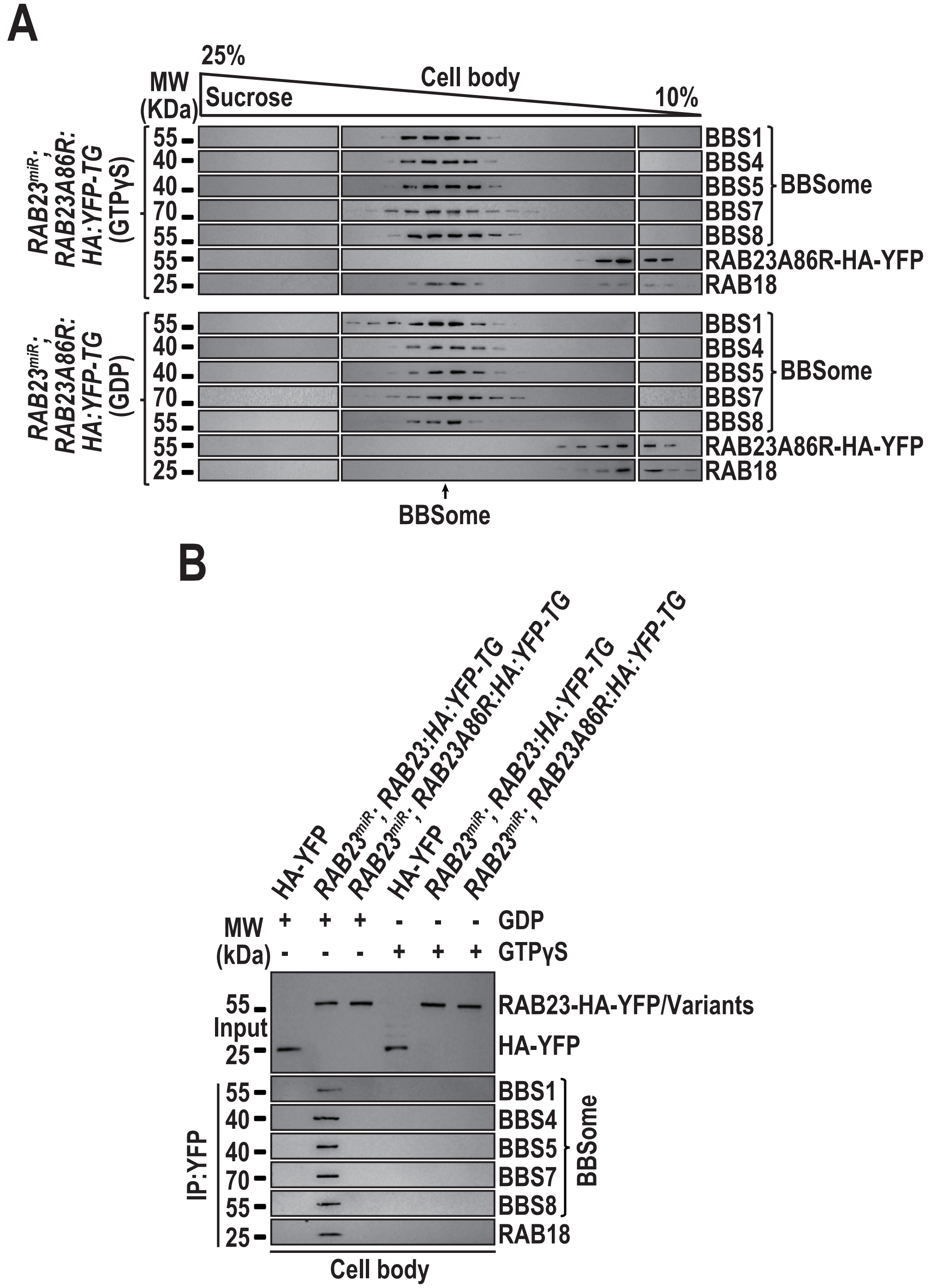

A
25%
Cell body
MW (KDa)
Sucrose
10%
RAB23miR; RAB23A86R: HA:YFP-TG (GTPγS)
55
40
40
70
55
55
25
BBS1
BBS4
BBS5
BBS7
BBS8
BBSome
RAB23A86R-HA-YFP
RAB18
RAB23miR; RAB23A86R: HA:YFP-TG (GDP)
BBSome
B
HA-YFP
RAB23miR; RAB23:HA:YFP-TG
RAB23miR; RAB23A86R:HA:YFP-TG
MW (kDa)
GDP
GTPγS
Input
RAB23-HA-YFP/Variants
HA-YFP
IP:YFP
Cell body
